# RabbitVar: ultra-fast and accurate somatic small-variant calling on multi-core architectures

**DOI:** 10.1101/2023.01.06.522980

**Authors:** Hao Zhang, Honglei Song, Zekun Yin, Qixin Chang, Yanjie Wei, Beifang Niu, Bertil Schmidt, Weiguo Liu

## Abstract

The continuous development of next-generation sequencing (NGS) technology has led to extensive and frequent use of genomic analysis in cancer research. The associated production of large-scale NGS datasets establishes the need for high-precision somatic variant calling methods that are highly optimized on commonly used hardware platforms. We present RabbitVar (https://github.com/LeiHaoa/RabbitVar), a scalable variant caller that can detect small somatic variants from paired tumor/normal NGS data on modern multi-core CPUs. Our approach combines candidate-finding and machine-learning-based filtering strategies with optimized data structures and multi-threading to achieve both high accuracy and efficiency. We have compared the performance of RabbitVar to leading state-of-the-art callers (Strelka2, Mutect2, NeuSomatic, VarDict, VarScan2) on real-world HCC1395 breast cancer datasets under different sequencing conditions and contamination rates. The evaluation results demonstrate that RabbitVar achieves highly competitive F1-scores when calling SNVs. Moreover, when calling the more challenging indel variants, it consistently achieves the highest F1-scores. RabbitVar is able to process a paired tumor and normal whole human genome sequencing datasets with 80x depth in less than 20 minutes on a 48-core workstation outperforming all other tested variant callers in terms of efficiency.

## 1 Main

With the advancements of next-generation sequencing (NGS) technology, personalized medicine has become feasible. In particular, the identification of somatic mutations is essential for modern cancer treatment with examples including the development of individualized therapeutic cancer vaccines [10]. This task relies on sequencing of a patient’s tumour and healthy tissue. Cancer-specific variations can then be detected by comparing these two NGS datsets.

As a consequence, a number of corresponding algorithms for somatic variant calling from paired tumor and normal sequencing data have been proposed. Examples of state-of-the-art tools include MuTect2 [1], Strelka2 [6], VarScan2 [8], VarDict [9], and NeuSomatic [11]. Existing methods can be distinguished by their algorithmic approach (e.g. statistical-based or machine learning-based) and supported mutation classes (such as the ability to detect single nucleotide variants (SNVs), insertions and deletions (indels), or fusion genes).

Identification of SNVs can in many cases be performed with high accuracy while highly accurate detection of indels is still challenging [7]. Furthermore, tumor heterogeneity leads to different purity and noise levels in the sequenced samples which makes somatic variation calling more complex than detection of germline mutations. Variant calling is often a bottleneck in typical processing pipelines [5]. Thus, delivering highly accurate results within an acceptable runtime is an additional important consideration in practical personalized cancer treatment. However, most existing somatic variation calling tools do not fully exploit the compute power of modern multi-core architectures due to sub-optimal implementations, which in turn leads to long runtimes on commonly used hardware platforms. Thus, an efficient yet accurate variant caller that is highly optimized for modern workstations without the need of special-purpose hardware is of high importance to clinical practice and biological research.

To address this need, we propose Rabbit-Var – an efficient and scalable tool for highly accurate calling of somatic SNVs and indels on typical multi-core CPUs. RabbitVar features a heuristic-based calling method and a subsequent machine-learning-based filtering strategy. In order to achieve both computational efficiency and high sensitivity, we first implement a highly optimized pipeline to identify candidate variants based on the approach proposed in VarDict [9]. SNVs, indels, multiple-nucleotide variants (MNVs), and complex variants are called simultaneously. We then reduce false positives introduced by sequencing truncation through a series of statistical tests and a local realignment strategy.

At this stage, the heuristic-based calling method typically detects more true variants compared to other callers, but at the same time, also introduces more false positives [2], which in turn reduces precision. This feature is particularly pronounced when calling mutations with low VAF (variant allele frequency), as shown in the supplement Fig. S5. To improve the precision, we propose a new XGBoost-based method to further filter out false positives. When calling candidates, we generate statistical information about each variant. The XGBoost model relies on statistical information (including amongst others variant allele frequency, mapping quality, and strand bias odd ratio) introduced in the sequencing, mapping, or candidate calling steps.

In addition, RabbitVar has also been highly optimized by featuring multi-threading, a high-performance memory allocator, vectorization, and efficient data structures on modern multi-core CPUs. The combination of these optimizations makes it both highly efficient and scalable. For high depth sequencing datasets, the runtime is linear with the sequencing depths, as shown in Fig. 2-(c).

To improve robustness, we use a series of datasets sequenced under different conditions (e.g. different sequencers, sample preparation, sequencing depth, purity, etc.) to train the XGBoost models for SNVs and indels, respectively. We use models trained on publicly available Spike-IN HCC1395BL datasets generated by BamSurgeon [3] and the real-world HCC1395 breast cancer datasets from SEQCII [4] project for accuracy assessment. The detailed information of both training and evaluation datasets are described in Section Method (see Table 1 and 2).

**Table 1:**
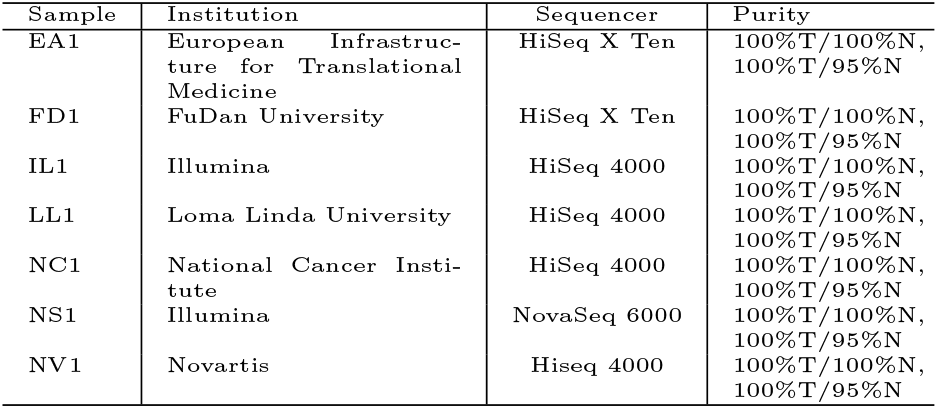
HCC1395/HCC1395BL data used for validation.

**Table 2:** The normal sample spiked by BAMSurgeon,

| Normal Sample | Tumor purity | Normal purity |
| --- | --- | --- |
| FD2 | 100% | 100%, 95% |
| FD3 | 100% | 100%, 95% |
| IL1 | 100% | 100%, 95% |
| IL2 | 100% | 100%, 95% |
| NS1 | 100% | 100%, 95% |
| NS2 | 100% | 100%, 95% |
| NS_1-5 | 100% | 100%, 95% |
| NS_5-9 | 100% | 100%, 95% |
| NV2 | 100% | 100%, 95% |
| NV3 | 100% | 100%, 95% |

To evaluate accuracy, we compare RabbitVar to five leading state-of-the-art tools: Strelka2 (v2.9.10), Mutect2 (v4.2.4.0), VarDict (v1.8.3), NeuSomatic(v0.2.1), and VarScan2 (v2.4.4) on different replicates sequenced at six sequencing centers with different purities and depths. The pre-processed data is publicly available in the Somatic Mutation Working Group of the SEQCII Consortium [14] website. The evaluation results are illustrated in Fig. 1 and 2. Note that all tested callers achieve high accuracy when calling SNVs. However, performance metrics are significantly lower for calling indels. Thus, improving the accuracy of indels is important, but also more challenging. In terms of overall accuracy, RabbitVar achieves significant and comprehensive improvement for calling indels under a variety of sequencing conditions, while clearly outperforming them in terms of efficiency. For example, RabbitVar achieves median speedups of 4.0x, 5.0x, 16.7x, 22.3x and 15.3x compared to Strelka2, Mutect2, VarDict, NeuSomatic and VarScan on the evaluation of runtime on the same hardware platform (48-core workstation) at various depths (10x, 30x, 50x, 80x, 200x, 300x).

**Fig. 1:**
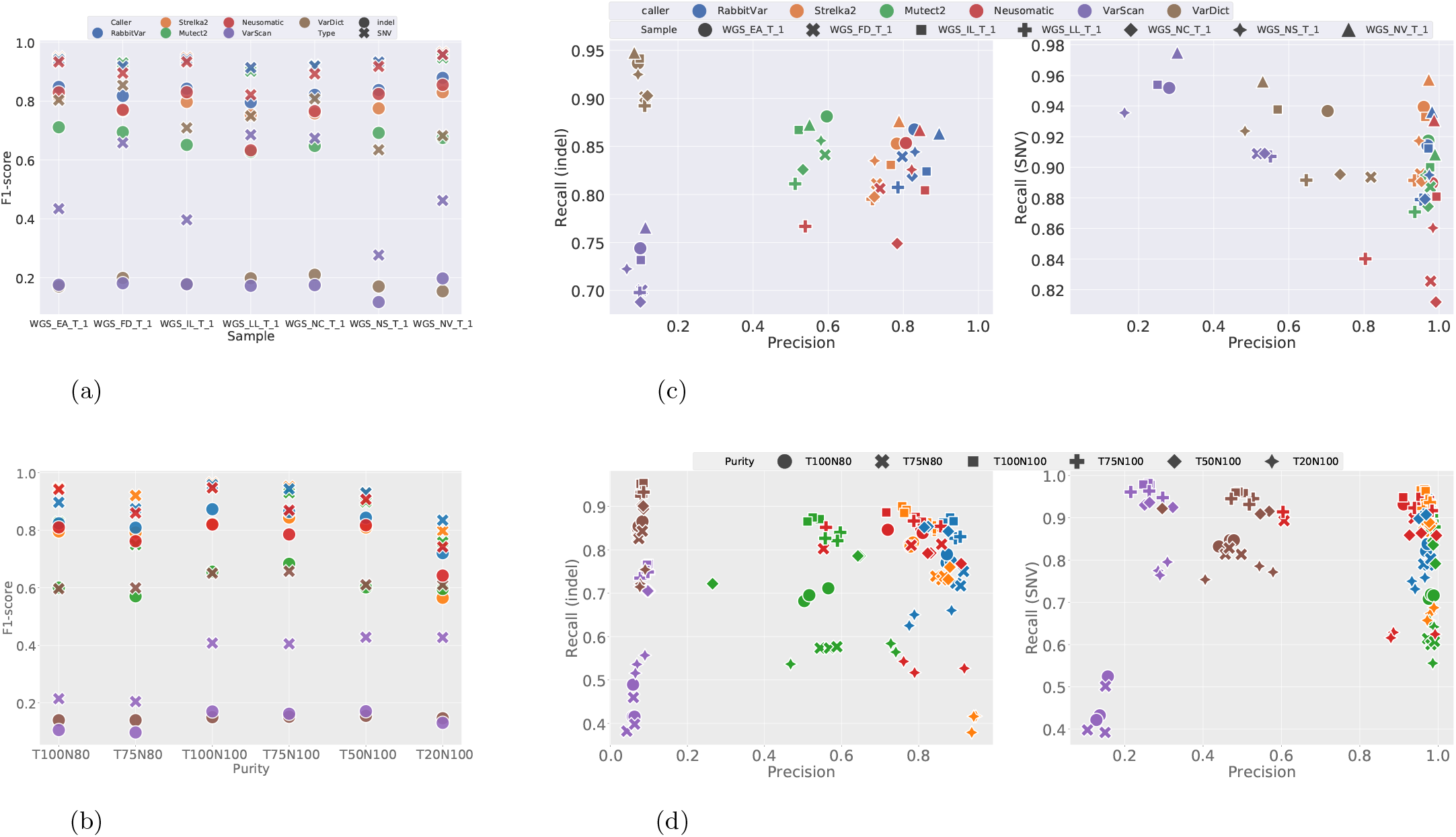
**(a)**. F1-score evaluation of SNV and indel calling under different sequencing conditions.**(b)**. F1-score evaluation of SNV and indel calling for different purity. **(c)**. Precision and recall evaluation of indel (left) and SNV (right) under different sequencing conditions. **(d)**. Precision and recall evaluation of indel (left) and SNV (right) for different purity.

**Fig. 2:**
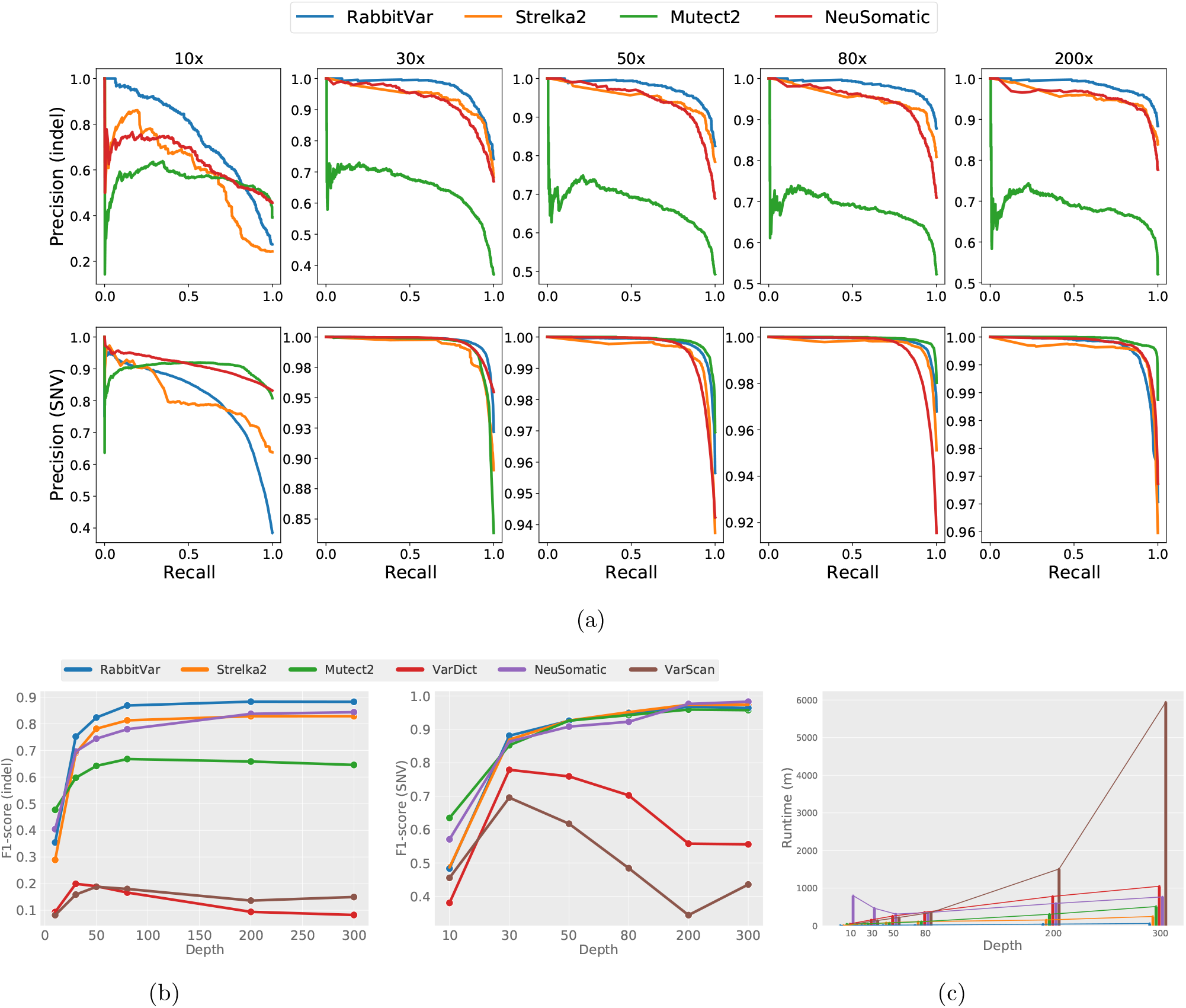
**(a)**. Scoring metrics used to generate curves were TLOD (Mutect2), RFV (RabbitVar), SomaticEVS (Strelka2), SCORE (NeuSomatic). Only PASS calls are used. Note that the scores for VarDict and VarScan are outside the axis limits due to low precision. **(b)**. F1-score of different caller with depth increase evaluated by hap.pyhttps://github.com/Illumina/hap.py. **(c)**. Runtime of different caller for increasing sequencing depths. Memory consumption is shown in Fig. S3 in the supplement. Experiments are conducted on a 48-core server with two AMD CPUs. When running NeuSomatic, we additionally use two NVIDIA GeForce RTX 3090 GPU in the calling step.

We have compared the performance of all callers on seven replicates under different sequencing conditions (see Table 1). As shown in Fig. 1-(a) and 1-(c), RabbitVar achieves the best performance for indel calling for all replicates with average F1-score improvements of 4.9%, 16.3%, 65.2%, 4.8% and 66.4% compared to Strelka2, Mutect2, VarDict, NeuSomatic and VarScan, respectively. Although VarDict and VarScan achieve the highest average recall, their overall accuracy is poor due to the low precision. The average F1-score for SNV calling of RabbitVar is also highly competitive: It is only slightly lower than Strelka2 (−0.42%), but higher than Mutect2 (+1.97%), VarDict (+23%), NeuSomatic (+2.5%) and VarScan (+41.9%).

Moreover, we have evaluated the accuracy of the six callers on impure tumor and normal samples. We have tested different combinations of tumor (100%, 75%, 50%, 20%) and normal (100% and 95%) purities. Three replicates of each combination are used to ensure reliability. As shown in Fig 1-(b), RabbitVar achieves higher F1-scores for indel calling than Strelka2, Mutect2, VariDict, NeuSomatic and VarScan with average improvements of 5.1%, 20.4%, 67.5%, 4.9% and 68.3% respectively. Particularly, it achieves significantly higher precision scores (2.6%, 30.8%, 78.9%, 8.7%, 79.1% average improvement compared to other callers).

Note that the impurity of normal samples makes all callers less sensitive, For example, at 75% tumor sample purity, a change in normal sample purity from 100% to 95% decreases the recall for indel calling of Strelka2, Mutect2, VarScan, RabbitVar, VarDict, and NeuSomatic by 12%, 25%, 33%, 10%, 10% and 5%, respectively. Especially, VarScan and Mutect2 are sensitive to the normal sample impurity. Although NeuSomatic has the lowest decrease in recall, RabbitVar still achieves the highest F1-score.

In terms of SNV calling, the average F1-score of RabbitVar is lower than Strelka2 (−1.2%), but still higher than Mutect2 (5.4%), VarDict (28.3%), NeuSomatic (2.8%) and VarScan (55.8%). Especially for tumor purity of 20%, it achieves the best F1-score on both indel and SNV calling task. Detailed results on different purity combinations are provided in the supplement.

Furthermore, we have evaluated accuracy using the HCC1395 dataset with varying sequencing depths of 10x, 30x, 50x, 80x, 200x and 300x. The precision-recall curves in Fig 2-(a) show that RabbitVar provides the best overall performance for indel calling under various sequencing depths, and achieves comparable performance to SNV calling. Fig 2-(b) shows that it also achieves a F1-score improvement of at least 4% compared to the second best caller when depth is over 30x for indel calling. RabbitVar, Strelka2 and Mutect2 are able to improve performance for increasing sequencing depths, but the F1-scores of VarDict and VarScan decrease when depth exceeds 30x due to higher amounts of false positives. Moreover, there is nearly no accuracy improvement for all callers when sequencing depth exceeds 200x. Mutect2 has the best F1-score for a low depth of 10x tumor/normal samples variant calling, however, this is not a common scenario in clinical research.

We also evaluated the performance on a synthetic data set NA24631.PACA [2], which spikes real pancreatic cancer (PACA) mutations into normal samples NA24631, using the same model as the above experiments. The result (see Table T8 and T9 in the supplement) show that RabbitVar achieves the second-best F1-score on both the indel and SNV variant calling tasks.

We have compared runtimes on a 48-core dual-socket AMD workstation and the optimal number of threads for each caller^1^. When calling WGS variants, all tools use the callable-regions generated by bcbio-nextgen (https://github.com/bcbio/bcbio-nextgen). The evaluation results show that for depth of 10-300x, RabbitVar can outperform all other tested state-of-the-art tools with speedups of at least 1.1x, 4.4x, 5.2x, 15.3x, and 5.9x compared to Strelka2, Mutect2, VarDict, NeuSomatic and VarScan. On average, it achieves speedups of 3.8x, 5.7x, 14.4x, 36.6x, and 30.0x respectively. See Fig. 2-(c) and supplement Table T7 for more details.

Note that the efficiency of some callers is sensitive to the calling regions. For example, testing the callers with all callable regions causes severe performance degradation of Strelka2 and VarDict compared to using only the high confident regions provided by the SEQCII project. However, RabbitVar exhibits a stable performance when using different region configurations, as shown in the supplement.

Our performance evaluation shows that RabbitVar can outperform leading state-of-the-art variant calling tools in terms of efficiency with a comparable or even higher accuracy. Thus, it can provide a high level of accuracy that can be delivered within an acceptable runtime on modern multi-core CPUs, e.g., typical human genome sequencing datasets with 80x depth can be processed in less than 20 minutes, which makes it an attractive tool for clinical cancer research.

## 2 Methods

### 2.1 RabbitVar Workflow

RabbitVar takes aligned sequencing reads (in BAM format) and a reference as input files and reports variant predictions as output. We adopt the VarDict pipeline and optimzed it with C++ to achieve better efficiency. The workflow is illustrated in Fig 3 and includes four main parts: (i) preprocessing, (ii) finding candidate variations, (iii) variation realignment, and (iv) filtration and formatting.

**Fig. 3:**
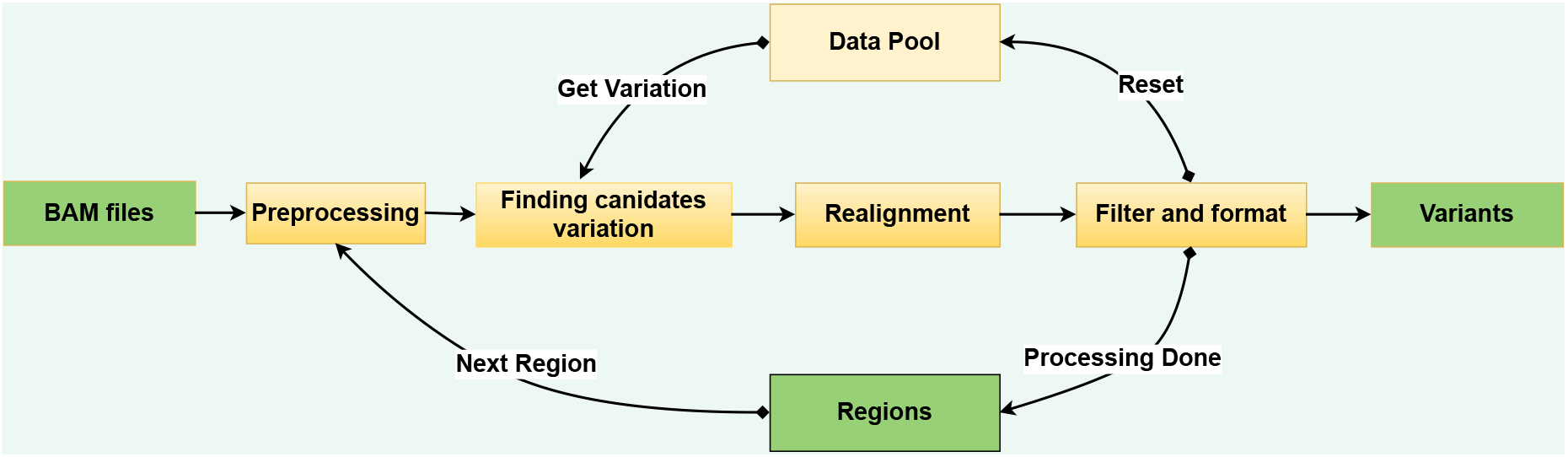
RabbitVar processing pipeline

#### Preprocessing

The preprocessing module receives target regions from a BED file (or the command line) and cleans reads. The primary purpose of preprocessing is to prepare the reference information according to the region information and to discard low-quality reads. Read quality is typically measured by average base quality scores, mapping quality scores, and the number of mismatches to the reference genome. For example, reads with low mapping quality or low average base quality, and duplicated reads are removed [13]. User-specified filtration rules are also supported.

#### Finding candidate variations

In this step, RabbitVar parses records and modifies the original CIGAR information provided in the input BAM files to facilitate subsequent operations. We process matches/mismatches (M), insertion (I), deletion (D), softclip (S) separately according to the modified CIGAR information. Preliminary candidate variants are identified by combining the sequence information of the sample and reference. We manipulate CIGAR arrays directly instead of converting them to *strings* in order to reduce runtime.

#### Local Realignment

It is challenging for read mappers to detect indels at the end of reads because gap-opening penalties used for alignment are typically larger than mismatch penalties. Thus, RabbitVar applies local realignment based on a heuristic strategy to reduce the number of false-positive variants and correct variant allele frequencies. First, soft-clip sequences starting/ending at the same reference position are collected to generate a consensus sequence. Second, we search forward to find consistent sequences to locate potential indels or add evidence to already-existing indels. The usage of local realignment makes our approach more accurate to calculate variant allele frequencies and more sensitive to low-frequency variants. Local realignment is also helpful for finding more complex variants. Reads supporting a complex variant are often misaligned or soft-clipped. Soft-clipping information that is close to each other in the 3’ end and 5’ end is collected. If the corresponding consensus sequence is ungapped matched with a mismatch. ≤ 3, it will be merged into the same variant.

#### Filtering and formatting

A number of basic hard filters are applied based on a series of heuristic conditions such as variant frequency, mean position, and mapping quality to remove potential false-positive variations. Subsequently, a pre-trained XGBoost-based filter is applied to further improve accuracy. Finally, variations are formatted and written to result files.

### 2.2 Performance Optimizations

Variant calling is often a bottleneck when dealing with large-scale sequencing data. We target our design for widely used multi-core CPU architectures. Our implementation features a number of key optimizations in order to take full advantage of modern hardware platforms:

- A carefully-tuned multi-threading strategy that balances the workload and reduces overheads.
- A data pool to remove redundant memory reallocations.
- Accelerated variant data manipulation using a highly optimized hash map.

#### Multi-threading

A straightforward approach to parallelize variation detection exploits the independence between different genomic regions. Thus, we base our solution on a load-balanced multi-threading method using C++ with OpenMP directives. Multiple threads work simultaneously: fetching regions from the region list specified in a BED file, performing the processing steps to detect variations, and finally writing variations to output files. To minimize load imbalance caused by different region sizes, we employ a dynamic scheduling strategy. In order to reduce the associated overhead of thread switching, we adapt set thread affinities according to the hardware configuration.

#### Data pool

When performing whole-genome analysis, worker threads allocate and free private intermediate memory for each task in duplicate fashion which in turn leads to memory reallocation redundancy. We have designed a data pool to avoid this overhead. Each worker thread initializes a data pool to store intermediate variation data. During execution, intermediate data is provided by the data pool and will be reset before handling the next region (see 3).

The initial size of the data pool needs to be carefully considered: large sizes may lead to an unnecessary high memory consumption while small sizes may lead to resizing overheads. Our experiments have shown that *memory pool size* = 1.2 * *average region size* is a good trade-off.

To avoid excessive memory consumption caused by large region sizes, we use an adaptive strategy. Our strategy does not miss any variant because we extend every region by 200-bps.

These optimizations allow RabbitVar to outperform other callers in terms of runtime efficiency. In terms of thread-level parallelism, it achieves near-linear thread scalability, which is shown in supplementary Figure S5.

### 2.3 XGBoost-based filter

To take full usage of the information generated by the first step, we train an XGBoost model to remove as many false positives as possible while retaining truth positive variants.

Before XGBoost-based filtering, several hard filters are applied to remove variants that are likely false positives. These standard filters include:

1. Low VAF variants (default 0.01, can be specified by user).
2. Mean quality (QM) ≤ 20.
3. Number of mismatch (NM) ≥ 6.
4. Variants labeled as “Germline.”

Furthermore, users can specify other filter parameters in the candidate set generation step.

To avoid overfitting due to the identical data distributions, we train filter models with synthetic data generated by BAMSurgeon and tested the accuracy with real-world HCC1395 and HCC1395BL samples.

There are about 92k SNV mutations and 22k indel mutations spiked in the normal sample reads where VAF is randomly selected from a beta distribution with the parameter *α* = 2, *β* = 5. The spiked-in normal samples are listed in Table 2. The trained sample covers different purity, library preparation protocol and depth to ensure robustness. For all samples, sequencing files are initially trimmed using Trimmomatic and then aligned with BWA-MEM (v0.7.15), followed by Picard MarkDuplicate. The reference genome version is GRCh38.d1.vd1 (https://gdc.cancer.gov/about-data/gdc-data-processing/gdc-reference-files). The utilized spiked-in normal samples will not be treated as validation datasets to ensure fairness. For each of the tumor-normal pairs, the impure normal by mixing 95% normal and 5% tumor are also used to train the models. Thus, there are 20 tumor-normal pair to be trained.

We provide scripts to create training datasets and train XGBoost models. Users need to specify: (1) candidate variant data generated by RabbitVar in the first step, (2) set of true variants (VCF file with the filter field marked as “PASS”) and (3) the type of data to be generated (SNV, indel).

In the training data generating step, the “chromosome name (CHR)”, “start position (POS)”, “reference allele (REF)”, and “alternative allele (ALT)” fields are combined into one field (“CHR:POS:REF:ALT” format). If the field in the variant data also exists in the true set, the label of this variant data is set to 1, otherwise, it is set to 0. Finally, all the data are saved in CSV format to the output file as training data with the format of each record as: [feature1, feature2, …, feature*n*, label].

The filter model uses different sets of features for SNV and indel filter tasks. The SNV feature set contains information in tumor and normal samples (some important features: MeanPosition, MismatchNumber, MappingQuality, Strand-Bias, Pvalue, and OddRatio), and extra information about the variant (such as MSI, MSINT, TumorNormalOddRatio). The indel feature set contains two additional features entries: VarType (Insertion, Deletion, MNV, Complex) and VarLabel (StrongSomatic, LikelySomatic, StrongLOH, LikelyLOH, Germline, and Adiff). None of the selected features are depth-related or sample-related in order to avoid sample or aligner-specific issues.

We use a grid search to find the best parameters for the XGBoost models and train the indel filter model with tree number = 1000, learning rate = 0.01, max depth = 20, min child weight = 16 (introduce the parameter). The SNV filter model uses tree number = 800, learning rate = 0.01, max depth = 25, min child weight = 18.

As shown in Figure S8 in the supplement, there are more false positive variants in the low-frequency interval. Thus, it is appropriate to adopt a segmented filtering strategy. We support both two-segmented scale filtering strategy, for example, a parameter with *“–scale 0*.*2:0*.*9:0*.*2”* means when VAF ≤ 0.2, the predict probability (*proba*) ≤ 0.9 will be filtered out, and when VAF > 0.2 the *proba ≤* 0.2 will be filtered out. Or a uniform scale filtering strategy, for example, *“–scale 0*.*5”* mean for all VAF, the variant will be filtered out when *proba ≤* 0.5. The default value is “0.2:0.9:0.2”.

### 2.4 Experimental Design

We use synthetic data to train the filter models and use real-world data for validation. Detailed information of the data sequenced under various conditions are listed in Table 1. For example, FD1 means the tissue processed and sequenced by FuDan University with the library preparation protocol #1 and Illumina HiSeq X Ten sequencer.

#### 1. Real world tumor-normal sample data

We use real-world tumor-normal data [4] provided by the Somatic Mutation Working Group of SEQC-II consortium. The consortium aims to develop guidelines for somatic mutation detection and provide golden-standard variant sets for wide-accepted paired tumor/normal reference samples/materials (HCC1395, HCC1395BL).

#### 2. Synthetic data

Synthetic data is created using BamSurgeon to spike the variants into different normal samples. Variants are generated with a minimal variant frequency of 0.01 and a maximum variant frequency of 1.0. The distribution of mutation frequency is *β*-distribution with *α* = 2 and *β* = 5. The minimum number of reads per variant is set to BAM files provided by SEQC-II project [12]. Every sample contains different purity as listed by “Normal purity” in Table 2.

In addition, we also use the synthetic data NA24631.PACA provided by [2]. The real pancreatic cancer (PACA) mutations are spiked into normal samples from Genome in a Bottle [15]’s NA24631, and spike-in frequencies uniformly sampled between 0.5% and 50%.

RabbitVar is implemented in C++ and Python. All runtime experiments have been conducted on a Linux server with two 24-core AMD EPYC 7402 CPUs, 256GB DDR4, a Samsung 970 EVO Plus NVME SSD (2TB), and a raid HDD array (MegaRAID SAS-3 3108 FCH SATA, RAID 5) running Centos 8.3. We provide the pre-trained model based on using the synthetic datasets, which is used for all evaluations in this paper. We also provide a pre-trained model using both synthetic and real-world datasets.

## Footnotes

1 Due to the lack of native support of Mutect2 and VarDict for multi-threading, we use bcbio-nextgen to run them to get better performance on multi-core systems

