## Supplementary for "RabbitVar: ultra-fast and accurate somatic small-variant calling on multi-core architectures"

### RabbitVar Supplementary

Sep 2022

#### 1 Evaluation Results

##### 1.1 Evaluation results of multiple callers (Strelka, Mutect2, VarDict, RabbitVar, NeuSomatic and VarScan) on HCC1395 samples

Table T1: Evaluation INDEL results of multiple callers (Strelka, Mutect2, VarDict, RabbitVar, NeuSomatic, VarScan) on HCC1395 samples.

| Caller & Data | Recall | Precision | F1-Score |
|---|---|---|---|
| WGS_EA_T_1_Strelka2 | 0.852923 | 0.782609 | 0.816254 |
| WGS_EA_T_1_Mutect2 | 0.881231 | 0.596170 | 0.711199 |
| WGS_EA_T_1_VarDict | 0.936615 | 0.093460 | 0.169961 |
| WGS_EA_T_1_RabbitVar | 0.867692 | 0.829412 | 0.848120 |
| WGS_EA_T_1_Neusomatic | 0.853538 | 0.806864 | 0.829545 |
| WGS_EA_T_1_VarScan | 0.744000 | 0.099695 | 0.175829 |
| WGS_FD_T_1_Strelka2 | 0.811077 | 0.728982 | 0.767842 |
| WGS_FD_T_1_Mutect2 | 0.841231 | 0.591519 | 0.694614 |
| WGS_FD_T_1_VarDict | 0.901538 | 0.111875 | 0.199049 |
| WGS_FD_T_1_RabbitVar | 0.839385 | 0.797195 | 0.817746 |
| WGS_FD_T_1_Neusomatic | 0.806154 | 0.737613 | 0.770362 |
| WGS_FD_T_1_VarScan | 0.700308 | 0.103993 | 0.181095 |
| WGS_IL_T_1_Strelka2 | 0.830769 | 0.766610 | 0.797401 |
| WGS_IL_T_1_Mutect2 | 0.867077 | 0.521466 | 0.651260 |
| WGS_IL_T_1_VarDict | 0.941538 | 0.098329 | 0.178062 |
| WGS_IL_T_1_RabbitVar | 0.824000 | 0.861647 | 0.842403 |
| WGS_IL_T_1_Neusomatic | 0.804308 | 0.857612 | 0.830105 |
| WGS_IL_T_1_VarScan | 0.731692 | 0.101131 | 0.177701 |
| WGS_LL_T_1_Strelka2 | 0.795077 | 0.717778 | 0.754453 |
| WGS_LL_T_1_Mutect2 | 0.811077 | 0.512044 | 0.627769 |
| WGS_LL_T_1_VarDict | 0.892308 | 0.111418 | 0.198101 |
| WGS_LL_T_1_RabbitVar | 0.807385 | 0.785629 | 0.796358 |
| WGS_LL_T_1_Neusomatic | 0.766769 | 0.538927 | 0.632969 |
| | | | Continued on next page |

| Caller & Data | Recall | Precision | F1-Score |
|---|---|---|---|
| WGS_LL_T_1_VarScan | 0.697846 | 0.098352 | 0.172406 |
| WGS_NC_T_1_Strelka2 | 0.797538 | 0.722811 | 0.758338 |
| WGS_NC_T_1_Mutect2 | 0.825846 | 0.532751 | 0.647683 |
| WGS_NC_T_1_VarDict | 0.902769 | 0.118853 | 0.210052 |
| WGS_NC_T_1_RabbitVar | 0.819077 | 0.823639 | 0.821351 |
| WGS_NC_T_1_Neusomatic | 0.748923 | 0.783645 | 0.765890 |
| WGS_NC_T_1_VarScan | 0.688000 | 0.100027 | 0.174660 |
| WGS_NS_T_1_Strelka2 | 0.835077 | 0.723733 | 0.775429 |
| WGS_NS_T_1_Mutect2 | 0.856000 | 0.580793 | 0.692040 |
| WGS_NS_T_1_VarDict | 0.924923 | 0.093732 | 0.170215 |
| WGS_NS_T_1_RabbitVar | 0.844308 | 0.831012 | 0.837607 |
| WGS_NS_T_1_Neusomatic | 0.825846 | 0.822304 | 0.824071 |
| WGS_NS_T_1_VarScan | 0.722462 | 0.063818 | 0.117277 |
| WGS_NV_T_1_Strelka2 | 0.876308 | 0.788483 | 0.830079 |
| WGS_NV_T_1_Mutect2 | 0.872615 | 0.550252 | 0.674917 |
| WGS_NV_T_1_VarDict | 0.947692 | 0.083705 | 0.153823 |
| WGS_NV_T_1_RabbitVar | 0.863385 | 0.895341 | 0.879073 |
| WGS_NV_T_1_Neusomatic | 0.867077 | 0.843713 | 0.855235 |
| WGS_NV_T_1_VarScan | 0.765538 | 0.113390 | 0.197523 |

Table T2: Evaluation SNV results of multiple callers (Strelka, Mutect2, VarDict, RabbitVar, Neusomatic, VarScan) on HCC1395 samples.

| Caller & Data | Recall | Precision | F1-Score |
|---|---|---|---|
| WGS_EA_T_1_Strelka2 | 0.939412 | 0.959082 | 0.949145 |
| WGS_EA_T_1_Mutect2 | 0.917383 | 0.971307 | 0.943575 |
| WGS_EA_T_1_VarDict | 0.936751 | 0.703231 | 0.803365 |
| WGS_EA_T_1_rabbitvar | 0.914239 | 0.969072 | 0.940857 |
| WGS_EA_T_1_Neusomatic | 0.889649 | 0.981239 | 0.933202 |
| WGS_EA_T_1_VarScan | 0.951834 | 0.281477 | 0.434471 |
| WGS_FD_T_1_Strelka2 | 0.895607 | 0.950241 | 0.922115 |
| WGS_FD_T_1_Mutect2 | 0.887114 | 0.976450 | 0.929641 |
| WGS_FD_T_1_VarDict | 0.893503 | 0.818380 | 0.854293 |
| WGS_FD_T_1_rabbitvar | 0.879028 | 0.960340 | 0.917887 |
| WGS_FD_T_1_Neusomatic | 0.825538 | 0.977488 | 0.895110 |
| WGS_FD_T_1_VarScan | 0.908865 | 0.516027 | 0.658294 |
| WGS_IL_T_1_Strelka2 | 0.932923 | 0.963276 | 0.947856 |
| WGS_IL_T_1_Mutect2 | 0.899992 | 0.976054 | 0.936481 |
| WGS_IL_T_1_VarDict | 0.937765 | 0.570160 | 0.709155 |
| WGS_IL_T_1_rabbitvar | 0.912287 | 0.971755 | 0.941083 |
| WGS_IL_T_1_Neusomatic | 0.880777 | 0.992856 | 0.933464 |
| WGS_IL_T_1_VarScan | 0.953913 | 0.250394 | 0.396667 |
| WGS_LL_T_1_Strelka2 | 0.891399 | 0.934341 | 0.912365 |

Continued on next page

| Caller & Data | Recall | Precision | F1-Score |
|---|---|---|---|
| WGS_LL_T_1_Mutect2 | 0.870789 | 0.935279 | 0.901883 |
| WGS_LL_T_1_VarDict | 0.891551 | 0.646346 | 0.749401 |
| WGS_LL_T_1_rabbitvar | 0.878850 | 0.951529 | 0.913747 |
| WGS_LL_T_1_Neusomatic | 0.840140 | 0.803710 | 0.821522 |
| WGS_LL_T_1_VarScan | 0.907065 | 0.550884 | 0.685466 |
| WGS_NC_T_1_Strelka2 | 0.890714 | 0.953306 | 0.920948 |
| WGS_NC_T_1_Mutect2 | 0.874363 | 0.970484 | 0.919919 |
| WGS_NC_T_1_VarDict | 0.895303 | 0.736461 | 0.808151 |
| WGS_NC_T_1_rabbitvar | 0.879154 | 0.962505 | 0.918943 |
| WGS_NC_T_1_Neusomatic | 0.811900 | 0.991395 | 0.892714 |
| WGS_NC_T_1_VarScan | 0.909017 | 0.535610 | 0.674054 |
| WGS_NS_T_1_Strelka2 | 0.917256 | 0.946109 | 0.931459 |
| WGS_NS_T_1_Mutect2 | 0.894922 | 0.962616 | 0.927535 |
| WGS_NS_T_1_VarDict | 0.923670 | 0.483172 | 0.634459 |
| WGS_NS_T_1_rabbitvar | 0.894948 | 0.973849 | 0.932733 |
| WGS_NS_T_1_Neusomatic | 0.860344 | 0.983682 | 0.917888 |
| WGS_NS_T_1_varscan | 0.935610 | 0.162499 | 0.276905 |
| WGS_NV_T_1_Strelka2 | 0.957259 | 0.972495 | 0.964817 |
| WGS_NV_T_1_Mutect2 | 0.908409 | 0.989043 | 0.947012 |
| WGS_NV_T_1_VarDict | 0.956068 | 0.530563 | 0.682421 |
| WGS_NV_T_1_rabbitvar | 0.936244 | 0.981242 | 0.958215 |
| WGS_NV_T_1_Neusomatic | 0.930743 | 0.986671 | 0.957891 |
| WGS_NV_T_1_varscan | 0.974928 | 0.303019 | 0.462338 |

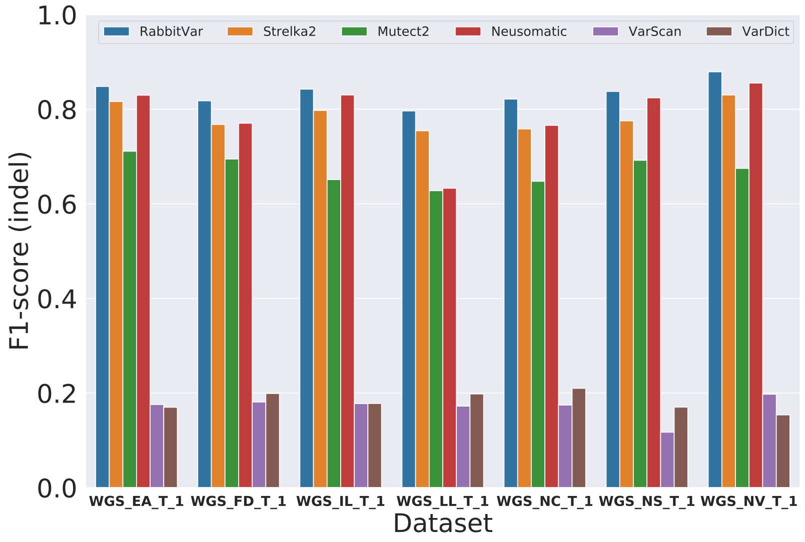


Figure S1: Indel calling accuracy on eight different samples.

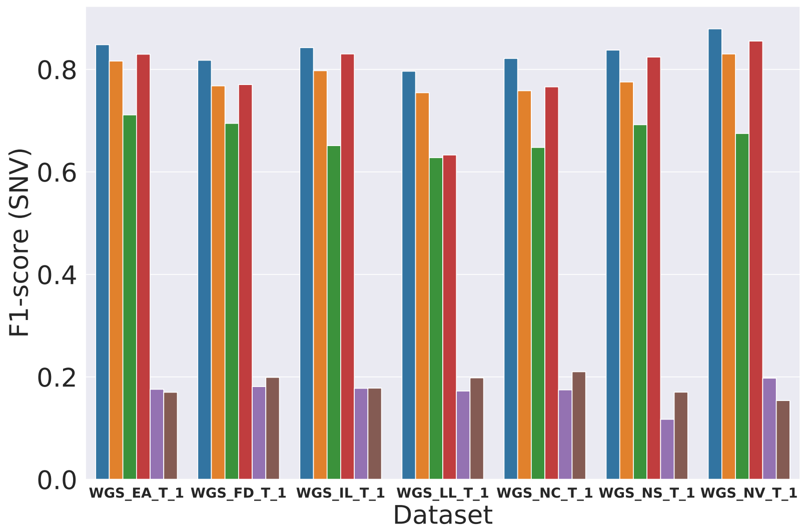


Figure S2: SNV calling accuracy on eight different samples.

#### 1.2 Evaluation results of different purity on multiple callers (Strelka, Mutect2, VarDict, RabbitVar, NeuSomatic, VarScan)

Table T3: SNVs calling result of different purity on Strelka2, Mutect2, VarDict, RabbitVar, Neusomatic and VarScan

| Caller | Purity | Recall | Precision | F1-Score |
|---|---|---|---|---|
| Strelka2 | T100N95_1 | 0.927548 | 0.955202 | 0.941172 |
| Mutect2 | T100N95_1 | 0.708394 | 0.976176 | 0.821001 |
| VarDict | T100N95_1 | 0.832231 | 0.440368 | 0.575968 |
| RabbitVar | T100N95_1 | 0.821254 | 0.969186 | 0.889108 |
| Neusomatic | T100N95_1 | 0.931883 | 0.911865 | 0.921765 |
| VarScan | T100N95_1 | 0.524552 | 0.156851 | 0.241492 |
| Strelka2 | T100N95_2 | 0.929374 | 0.968945 | 0.948747 |
| Mutect2 | T100N95_2 | 0.718686 | 0.981954 | 0.829942 |
| VarDict | T100N95_2 | 0.847035 | 0.468501 | 0.603308 |
| RabbitVar | T100N95_2 | 0.838568 | 0.973513 | 0.901016 |
| Neusomatic | T100N95_2 | 0.939336 | 0.957344 | 0.948255 |
| VarScan | T100N95_2 | 0.432479 | 0.136136 | 0.207085 |
| Strelka2 | T100N95_3 | 0.927041 | 0.972554 | 0.949252 |
| Mutect2 | T100N95_3 | 0.716252 | 0.988524 | 0.830646 |
| VarDict | T100N95_3 | 0.846858 | 0.478452 | 0.611451 |
| RabbitVar | T100N95_3 | 0.837985 | 0.973123 | 0.900512 |
| Neusomatic | T100N95_3 | 0.934139 | 0.978829 | 0.955962 |
| VarScan | T100N95_3 | 0.421502 | 0.127591 | 0.195886 |
| Strelka2 | T75N95_1 | 0.880143 | 0.968344 | 0.922139 |
| Mutect2 | T75N95_1 | 0.614166 | 0.973011 | 0.753023 |
| VarDict | T75N95_1 | 0.814054 | 0.456694 | 0.585126 |
| RabbitVar | T75N95_1 | 0.787943 | 0.975060 | 0.871572 |
| Neusomatic | T75N95_1 | 0.898776 | 0.938582 | 0.918248 |
| VarScan | T75N95_1 | 0.501483 | 0.150803 | 0.231877 |
| Strelka2 | T75N95_2 | 0.867899 | 0.963906 | 0.913386 |
| Mutect2 | T75N95_2 | 0.599894 | 0.981746 | 0.744725 |
| VarDict | T75N95_2 | 0.813547 | 0.497334 | 0.617302 |
| RabbitVar | T75N95_2 | 0.790783 | 0.962986 | 0.868430 |
| Neusomatic | T75N95_2 | 0.893249 | 0.606691 | 0.722597 |
| VarScan | T75N95_2 | 0.391969 | 0.149798 | 0.216758 |
| Strelka2 | T75N95_3 | 0.877684 | 0.980654 | 0.926316 |
| Mutect2 | T75N95_3 | 0.607549 | 0.990494 | 0.753139 |
| VarDict | T75N95_3 | 0.828682 | 0.466439 | 0.596901 |
| RabbitVar | T75N95_3 | 0.803356 | 0.980902 | 0.883296 |
| Neusomatic | T75N95_3 | 0.897584 | 0.984896 | 0.939215 |
| VarScan | T75N95_3 | 0.397926 | 0.104783 | 0.165885 |

Continued on next page

| Caller | Purity | Recall | Precision | F1-Score |
|---|---|---|---|---|
| Strelka2 | T100N100_1 | 0.965042 | 0.955786 | 0.960392 |
| Mutect2 | T100N100_1 | 0.915760 | 0.985003 | 0.949120 |
| VarDict | T100N100_1 | 0.958501 | 0.503764 | 0.660425 |
| RabbitVar | T100N100_1 | 0.943291 | 0.968153 | 0.955560 |
| Neusomatic | T100N100_1 | 0.948716 | 0.910759 | 0.929350 |
| VarScan | T100N100_1 | 0.977869 | 0.265334 | 0.417409 |
| Strelka2 | T100N100_2 | 0.967222 | 0.964874 | 0.966046 |
| Mutect2 | T100N100_2 | 0.917053 | 0.983711 | 0.949213 |
| VarDict | T100N100_2 | 0.962076 | 0.490171 | 0.649451 |
| RabbitVar | T100N100_2 | 0.946485 | 0.974550 | 0.960313 |
| Neusomatic | T100N100_2 | 0.949679 | 0.954713 | 0.952190 |
| VarScan | T100N100_2 | 0.982381 | 0.260320 | 0.411577 |
| Strelka2 | T100N100_3 | 0.963293 | 0.968152 | 0.965716 |
| Mutect2 | T100N100_3 | 0.910259 | 0.989283 | 0.948127 |
| VarDict | T100N100_3 | 0.959820 | 0.483100 | 0.642709 |
| RabbitVar | T100N100_3 | 0.942429 | 0.976261 | 0.959047 |
| Neusomatic | T100N100_3 | 0.942049 | 0.979003 | 0.960171 |
| VarScan | T100N100_3 | 0.978883 | 0.247475 | 0.395071 |
| Strelka2 | T75N100_1 | 0.940097 | 0.971625 | 0.955601 |
| Mutect2 | T75N100_1 | 0.892184 | 0.985495 | 0.936521 |
| VarDict | T75N100_1 | 0.945471 | 0.527771 | 0.677407 |
| RabbitVar | T75N100_1 | 0.919614 | 0.976500 | 0.947204 |
| Neusomatic | T75N100_1 | 0.923416 | 0.939517 | 0.931397 |
| VarScan | T75N100_1 | 0.963318 | 0.262469 | 0.412537 |
| Strelka2 | T75N100_2 | 0.922428 | 0.962212 | 0.941900 |
| Mutect2 | T75N100_2 | 0.869876 | 0.984394 | 0.923599 |
| VarDict | T75N100_2 | 0.931680 | 0.517947 | 0.665773 |
| RabbitVar | T75N100_2 | 0.907724 | 0.964550 | 0.935275 |
| Neusomatic | T75N100_2 | 0.914113 | 0.603983 | 0.727370 |
| VarScan | T75N100_2 | 0.947727 | 0.296885 | 0.452135 |
| Strelka2 | T75N100_3 | 0.936168 | 0.979159 | 0.957181 |
| Mutect2 | T75N100_3 | 0.878444 | 0.990878 | 0.931280 |
| VarDict | T75N100_3 | 0.945294 | 0.471899 | 0.629531 |
| RabbitVar | T75N100_3 | 0.915532 | 0.983497 | 0.948298 |
| Neusomatic | T75N100_3 | 0.917003 | 0.984219 | 0.949423 |
| VarScan | T75N100_3 | 0.961163 | 0.215266 | 0.351752 |
| Strelka2 | T50N100_1 | 0.886937 | 0.983085 | 0.932539 |
| Mutect2 | T50N100_1 | 0.845337 | 0.986860 | 0.910633 |
| VarDict | T50N100_1 | 0.915482 | 0.568574 | 0.701482 |
| RabbitVar | T50N100_1 | 0.903136 | 0.956634 | 0.929115 |
| NeuSomatic | T50N100_1 | 0.863817 | 0.956921 | 0.907989 |
| VarScan | T50N100_1 | 0.930312 | 0.250539 | 0.394766 |
| Strelka2 | T50N100_2 | 0.880751 | 0.979587 | 0.927544 |
| Mutect2 | T50N100_2 | 0.835830 | 0.987451 | 0.905337 |

Continued on next page

| Caller | Purity | Recall | Precision | F1-Score |
|---|---|---|---|---|
| VarDict | T50N100_2 | 0.909195 | 0.545957 | 0.682240 |
| RabbitVar | T50N100_2 | 0.898142 | 0.951012 | 0.923821 |
| NeuSomatic | T50N100_2 | 0.858823 | 0.926819 | 0.891526 |
| VarScan | T50N100_2 | 0.924734 | 0.322689 | 0.478428 |
| Strelka2 | T50N100_3 | 0.889447 | 0.977108 | 0.931219 |
| Mutect2 | T50N100_3 | 0.791518 | 0.993161 | 0.880948 |
| VarDict | T50N100_3 | 0.921971 | 0.295678 | 0.447759 |
| RabbitVar | T50N100_3 | 0.907344 | 0.969395 | 0.937344 |
| NeuSomatic | T50N100_3 | 0.858189 | 0.995618 | 0.921810 |
| VarScan | T50N100_3 | 0.936294 | 0.263421 | 0.411164 |
| Strelka2 | T20N100_1 | 0.673359 | 0.980075 | 0.798269 |
| Mutect2 | T20N100_1 | 0.656096 | 0.978414 | 0.785475 |
| VarDict | T20N100_1 | 0.771567 | 0.578543 | 0.661257 |
| RabbitVar | T20N100_1 | 0.749867 | 0.932917 | 0.831436 |
| NeuSomatic | T20N100_1 | 0.628894 | 0.885779 | 0.735553 |
| VarScan | T20N100_1 | 0.775015 | 0.285107 | 0.416862 |
| Strelka2 | T20N100_2 | 0.687302 | 0.989164 | 0.811057 |
| Mutect2 | T20N100_2 | 0.641874 | 0.989565 | 0.778670 |
| VarDict | T20N100_2 | 0.785586 | 0.543724 | 0.642652 |
| RabbitVar | T20N100_2 | 0.758461 | 0.966189 | 0.849815 |
| NeuSomatic | T20N100_2 | 0.624509 | 0.992266 | 0.766562 |
| VarScan | T20N100_2 | 0.795498 | 0.308919 | 0.445021 |
| Strelka2 | T20N100_3 | 0.657363 | 0.972218 | 0.784374 |
| Mutect2 | T20N100_3 | 0.555885 | 0.986326 | 0.711036 |
| VarDict | T20N100_3 | 0.753872 | 0.405304 | 0.527181 |
| RabbitVar | T20N100_3 | 0.731488 | 0.941129 | 0.823170 |
| NeuSomatic | T20N100_3 | 0.615915 | 0.879302 | 0.724410 |
| VarScan | T20N100_3 | 0.764139 | 0.289784 | 0.420211 |

Table T4: Indel calling result of different purity on Strelka2, Mutect2, VarDict, RabbitVar, Neusomatic and VarScan

| Caller | Purity | Recall | Precision | F1-Score |
|---|---|---|---|---|
| Strelka2 | T100N95_1 | 0.812308 | 0.776928 | 0.794224 |
| Mutect2 | T100N95_1 | 0.681846 | 0.503179 | 0.579044 |
| VarDict | T100N95_1 | 0.849231 | 0.071722 | 0.132273 |
| RabbitVar | T100N95_1 | 0.769846 | 0.872385 | 0.817914 |
| Neusomatic | T100N95_1 | 0.846154 | 0.719895 | 0.777935 |
| VarScan | T100N95_1 | 0.489231 | 0.059130 | 0.105508 |
| Strelka2 | T100N95_2 | 0.816615 | 0.785207 | 0.800603 |
| Mutect2 | T100N95_2 | 0.695385 | 0.516217 | 0.592554 |
| VarDict | T100N95_2 | 0.854154 | 0.073517 | 0.135382 |
| RabbitVar | T100N95_2 | 0.772308 | 0.883803 | 0.824302 |

Continued on next page

| Caller | Purity | Recall | Precision | F1-Score |
|---|---|---|---|---|
| Neusomatic | T100N95_2 | 0.849846 | 0.806659 | 0.827690 |
| VarScan | T100N95_2 | 0.415385 | 0.058650 | 0.102787 |
| Strelka2 | T100N95_3 | 0.807385 | 0.778635 | 0.792749 |
| Mutect2 | T100N95_3 | 0.711385 | 0.565558 | 0.630144 |
| VarDict | T100N95_3 | 0.865231 | 0.083526 | 0.152346 |
| RabbitVar | T100N95_3 | 0.789538 | 0.873383 | 0.829347 |
| Neusomatic | T100N95_3 | 0.838769 | 0.810345 | 0.824312 |
| VarScan | T100N95_3 | 0.415385 | 0.062808 | 0.109117 |
| Strelka2 | T75N95_1 | 0.739077 | 0.844585 | 0.788316 |
| Mutect2 | T75N95_1 | 0.573538 | 0.544393 | 0.558586 |
| VarDict | T75N95_1 | 0.827692 | 0.072099 | 0.132643 |
| RabbitVar | T75N95_1 | 0.723077 | 0.896262 | 0.800409 |
| Neusomatic | T75N95_1 | 0.810462 | 0.780676 | 0.795290 |
| VarScan | T75N95_1 | 0.459692 | 0.060024 | 0.106183 |
| Strelka2 | T75N95_2 | 0.731077 | 0.860246 | 0.790419 |
| Mutect2 | T75N95_2 | 0.573538 | 0.568639 | 0.571078 |
| VarDict | T75N95_2 | 0.825846 | 0.074123 | 0.136036 |
| RabbitVar | T75N95_2 | 0.716923 | 0.908736 | 0.801514 |
| Neusomatic | T75N95_2 | 0.802462 | 0.553011 | 0.654783 |
| VarScan | T75N95_2 | 0.382154 | 0.042899 | 0.077138 |
| Strelka2 | T75N95_3 | 0.739077 | 0.867148 | 0.798007 |
| Mutect2 | T75N95_3 | 0.576615 | 0.587829 | 0.582168 |
| VarDict | T75N95_3 | 0.843077 | 0.083166 | 0.151398 |
| RabbitVar | T75N95_3 | 0.750154 | 0.916541 | 0.825042 |
| Neusomatic | T75N95_3 | 0.812923 | 0.859466 | 0.835547 |
| VarScan | T75N95_3 | 0.398154 | 0.063153 | 0.109014 |
| Strelka2 | T100N100_1 | 0.900308 | 0.757246 | 0.822603 |
| Mutect2 | T100N100_1 | 0.870769 | 0.543395 | 0.669189 |
| VarDict | T100N100_1 | 0.947692 | 0.080175 | 0.147842 |
| RabbitVar | T100N100_1 | 0.874462 | 0.881514 | 0.877973 |
| Neusomatic | T100N100_1 | 0.886154 | 0.716062 | 0.792079 |
| VarScan | T100N100_1 | 0.764308 | 0.092929 | 0.165710 |
| Strelka2 | T100N100_2 | 0.891692 | 0.768700 | 0.825641 |
| Mutect2 | T100N100_2 | 0.874462 | 0.525129 | 0.656199 |
| VarDict | T100N100_2 | 0.952000 | 0.078736 | 0.145443 |
| RabbitVar | T100N100_2 | 0.865846 | 0.890506 | 0.878003 |
| Neusomatic | T100N100_2 | 0.875692 | 0.792316 | 0.831920 |
| VarScan | T100N100_2 | 0.767385 | 0.099268 | 0.175795 |
| Strelka2 | T100N100_3 | 0.884923 | 0.762460 | 0.819140 |
| Mutect2 | T100N100_3 | 0.865231 | 0.511459 | 0.642890 |
| VarDict | T100N100_3 | 0.953846 | 0.086236 | 0.158171 |
| RabbitVar | T100N100_3 | 0.862769 | 0.863832 | 0.863300 |
| Neusomatic | T100N100_3 | 0.864615 | 0.808400 | 0.835563 |
| VarScan | T100N100_3 | 0.765538 | 0.095553 | 0.169899 |

Continued on next page

| Caller | Purity | Recall | Precision | F1-Score |
|---|---|---|---|---|
| Strelka2 | T75N100_1 | 0.858462 | 0.828385 | 0.843155 |
| Mutect2 | T75N100_1 | 0.840000 | 0.596591 | 0.697674 |
| VarDict | T75N100_1 | 0.932308 | 0.081614 | 0.150089 |
| RabbitVar | T75N100_1 | 0.829538 | 0.902880 | 0.864657 |
| Neusomatic | T75N100_1 | 0.866462 | 0.787472 | 0.825081 |
| VarScan | T75N100_1 | 0.747077 | 0.098236 | 0.173639 |
| Strelka2 | T75N100_2 | 0.846154 | 0.840465 | 0.843300 |
| Mutect2 | T75N100_2 | 0.820923 | 0.589223 | 0.686038 |
| VarDict | T75N100_2 | 0.923692 | 0.080212 | 0.147605 |
| RabbitVar | T75N100_2 | 0.826462 | 0.895333 | 0.859520 |
| Neusomatic | T75N100_2 | 0.851692 | 0.559644 | 0.675451 |
| VarScan | T75N100_2 | 0.734769 | 0.078075 | 0.141151 |
| Strelka2 | T75N100_3 | 0.851692 | 0.840826 | 0.846224 |
| Mutect2 | T75N100_3 | 0.826462 | 0.557724 | 0.666005 |
| VarDict | T75N100_3 | 0.932923 | 0.086505 | 0.158329 |
| RabbitVar | T75N100_3 | 0.830154 | 0.907806 | 0.867245 |
| Neusomatic | T75N100_3 | 0.854154 | 0.857319 | 0.855734 |
| VarScan | T75N100_3 | 0.748923 | 0.097485 | 0.172514 |
| Strelka2 | T50N100_1 | 0.759385 | 0.880799 | 0.815598 |
| Mutect2 | T50N100_1 | 0.785846 | 0.647566 | 0.710036 |
| VarDict | T50N100_1 | 0.903385 | 0.082975 | 0.151990 |
| RabbitVar | T50N100_1 | 0.853538 | 0.824614 | 0.838827 |
| NeuSomatic | T50N100_1 | 0.792615 | 0.831504 | 0.811594 |
| VarScan | T50N100_1 | 0.705846 | 0.094528 | 0.166727 |
| Strelka2 | T50N100_2 | 0.760000 | 0.880884 | 0.815989 |
| Mutect2 | T50N100_2 | 0.785846 | 0.642354 | 0.706892 |
| VarDict | T50N100_2 | 0.894154 | 0.086283 | 0.157379 |
| RabbitVar | T50N100_2 | 0.851692 | 0.814597 | 0.832732 |
| NeuSomatic | T50N100_2 | 0.792000 | 0.823417 | 0.807403 |
| VarScan | T50N100_2 | 0.705231 | 0.100579 | 0.176050 |
| Strelka2 | T50N100_3 | 0.731692 | 0.876844 | 0.797719 |
| Mutect2 | T50N100_3 | 0.721846 | 0.265385 | 0.388089 |
| VarDict | T50N100_3 | 0.900923 | 0.085504 | 0.156185 |
| RabbitVar | T50N100_3 | 0.842462 | 0.877002 | 0.859385 |
| NeuSomatic | T50N100_3 | 0.768000 | 0.910284 | 0.833111 |
| VarScan | T50N100_3 | 0.705231 | 0.097416 | 0.171185 |
| Strelka2 | T20N100_1 | 0.416615 | 0.948179 | 0.578880 |
| Mutect2 | T20N100_1 | 0.564308 | 0.740711 | 0.640587 |
| VarDict | T20N100_1 | 0.721846 | 0.077816 | 0.140487 |
| RabbitVar | T20N100_1 | 0.650462 | 0.788806 | 0.712985 |
| NeuSomatic | T20N100_1 | 0.542769 | 0.761001 | 0.633621 |
| VarScan | T20N100_1 | 0.536000 | 0.068968 | 0.122211 |
| Strelka2 | T20N100_2 | 0.416000 | 0.942817 | 0.577284 |
| Mutect2 | T20N100_2 | 0.584000 | 0.727761 | 0.648003 |

Continued on next page

| Caller | Purity | Recall | Precision | F1-Score |
|---|---|---|---|---|
| VarDict | T20N100_2 | 0.754462 | 0.090446 | 0.161528 |
| RabbitVar | T20N100_2 | 0.660308 | 0.885314 | 0.756433 |
| NeuSomatic | T20N100_2 | 0.526769 | 0.918455 | 0.669535 |
| VarScan | T20N100_2 | 0.556923 | 0.089853 | 0.154741 |
| Strelka2 | T20N100_3 | 0.379077 | 0.937595 | 0.539877 |
| Mutect2 | T20N100_3 | 0.536615 | 0.467811 | 0.499857 |
| VarDict | T20N100_3 | 0.714462 | 0.076689 | 0.138511 |
| RabbitVar | T20N100_3 | 0.625231 | 0.775573 | 0.692334 |
| NeuSomatic | T20N100_3 | 0.516923 | 0.789474 | 0.624768 |
| VarScan | T20N100_3 | 0.515692 | 0.065062 | 0.115546 |

#### 1.3 Evaluation results of different depth on Strelka2, Mutect2, VarDict, RabbitVar, NeuSomatic and VarScan

##### 1.3.1 Memory consumption

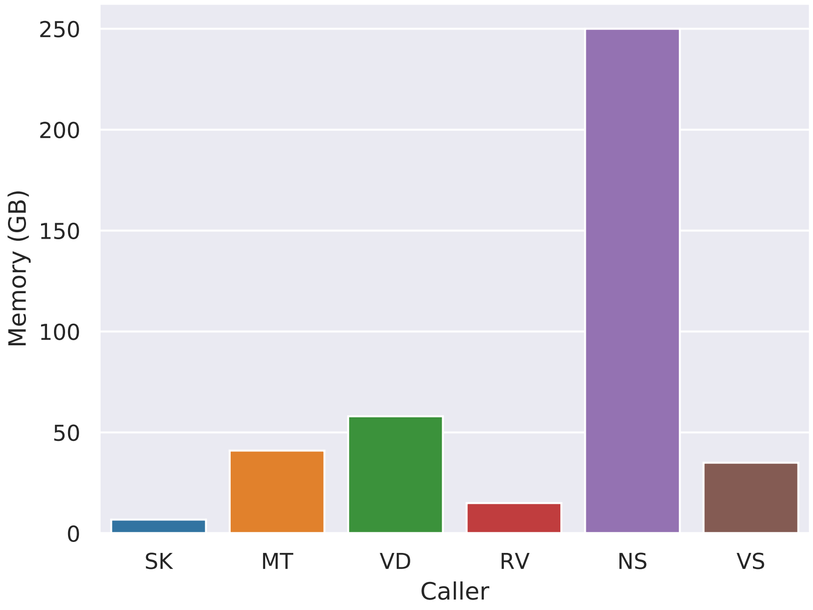


Figure S3: Memory consumption of Strelka2(SK), Mutect2(MT), VarDict(VD), RabbitVar(VD), NeuSomatic(NS), VarScan(VS) on 80x WGS dataset. We use the memusg (`https://github.com/ctsa/memusg`) to evaluate the peak rss memory consumption.

##### 1.3.2 Accuracy on different depth

Table T5: SNV calling result on different depth

| | | | | |
|---|---|---|---|---|
| 10x.snv | Strelka2 | 0.392780 | 0.639508 | 0.486659 |
| 10x.snv | Mutect2 | 0.523817 | 0.805355 | 0.634769 |
| 10x.snv | VarDict | 0.421730 | 0.346374 | 0.380356 |
| 10x.snv | RabbitVar | 0.649885 | 0.384769 | 0.483361 |
| 10x.snv | NeuSomatic | 0.434482 | 0.831587 | 0.570758 |
| 10x.snv | VarScan | 0.531599 | 0.398306 | 0.455399 |
| 30x.snv | Strelka2 | 0.848734 | 0.890876 | 0.869294 |
| 30x.snv | Mutect2 | 0.860344 | 0.844586 | 0.852392 |
| 30x.snv | VarDict | 0.847289 | 0.720805 | 0.778946 |
| 30x.snv | RabbitVar | 0.844272 | 0.921196 | 0.881058 |
| 30x.snv | NeuSomatic | 0.788374 | 0.954425 | 0.863489 |
| 30x.snv | VarScan | 0.865034 | 0.581718 | 0.695635 |
| 50x.snv | Strelka2 | 0.915786 | 0.937265 | 0.926401 |
| 50x.snv | Mutect2 | 0.885314 | 0.971054 | 0.926204 |
| 50x.snv | VarDict | 0.913910 | 0.649310 | 0.759216 |
| 50x.snv | RabbitVar | 0.897128 | 0.956330 | 0.925783 |
| 50x.snv | NeuSomatic | 0.876797 | 0.942143 | 0.908296 |
| 50x.snv | VarScan | 0.929348 | 0.462318 | 0.617468 |
| 80x.snv | Strelka2 | 0.952569 | 0.950978 | 0.951773 |
| 80x.snv | Mutect2 | 0.908028 | 0.980268 | 0.942767 |
| 80x.snv | VarDict | 0.949730 | 0.557293 | 0.702415 |
| 80x.snv | RabbitVar | 0.931224 | 0.967830 | 0.949174 |
| 80x.snv | NeuSomatic | 0.930261 | 0.915523 | 0.922833 |
| 80x.snv | VarScan | 0.967273 | 0.322926 | 0.484201 |
| 200x.snv | Strelka2 | 0.983294 | 0.964636 | 0.973875 |
| 200x.snv | Mutect2 | 0.932086 | 0.988307 | 0.959374 |
| 200x.snv | VarDict | 0.971278 | 0.391227 | 0.557781 |
| 200x.snv | RabbitVar | 0.965194 | 0.970384 | 0.967782 |
| 200x.snv | NeuSomatic | 0.978959 | 0.973505 | 0.976224 |
| 200x.snv | VarScan | 0.995031 | 0.207741 | 0.343721 |
| 300x.snv | Strelka2 | 0.985423 | 0.962202 | 0.973674 |
| 300x.snv | Mutect2 | 0.925571 | 0.991796 | 0.957540 |
| 300x.snv | VarDict | 0.972824 | 0.388990 | 0.555757 |
| 300x.snv | RabbitVar | 0.968185 | 0.959839 | 0.963994 |
| 300x.snv | NeuSomatic | 0.984612 | 0.981428 | 0.983017 |
| 300x.snv | VarScan | 0.997516 | 0.278520 | 0.435455 |

##### 1.3.3 Runtime evaluation

Table T6: indel calling result on different depth

| | | | | |
|---|---|---|---|---|
| 10x.indel | Strelka2 | 0.300308 | 0.278539 | 0.289014 |
| 10x.indel | Mutect2 | 0.416000 | 0.559140 | 0.477064 |
| 10x.indel | VarDict | 0.352615 | 0.053657 | 0.093140 |
| 10x.indel | RabbitVar | 0.508308 | 0.271979 | 0.354354 |
| 10x.indel | NeuSomatic | 0.364308 | 0.454336 | 0.404372 |
| 10x.indel | VarScan | 0.333538 | 0.046508 | 0.081633 |
| 30x.indel | Strelka2 | 0.719385 | 0.662698 | 0.689879 |
| 30x.indel | Mutect2 | 0.769231 | 0.488091 | 0.597229 |
| 30x.indel | VarDict | 0.832615 | 0.113127 | 0.199190 |
| 30x.indel | RabbitVar | 0.764923 | 0.739441 | 0.751966 |
| 30x.indel | NeuSomatic | 0.725538 | 0.667989 | 0.695575 |
| 30x.indel | VarScan | 0.639385 | 0.090592 | 0.158699 |
| 50x.indel | Strelka2 | 0.820308 | 0.745943 | 0.781360 |
| 50x.indel | Mutect2 | 0.814769 | 0.529388 | 0.641784 |
| 50x.indel | VarDict | 0.903385 | 0.106292 | 0.190205 |
| 50x.indel | RabbitVar | 0.824615 | 0.822086 | 0.823349 |
| 50x.indel | NeuSomatic | 0.812923 | 0.685877 | 0.744016 |
| 50x.indel | VarScan | 0.719385 | 0.108151 | 0.188033 |
| 80x.indel | Strelka2 | 0.880000 | 0.755016 | 0.812731 |
| 80x.indel | Mutect2 | 0.858462 | 0.546202 | 0.667624 |
| 80x.indel | VarDict | 0.938462 | 0.091268 | 0.166358 |
| 80x.indel | RabbitVar | 0.861538 | 0.875547 | 0.868486 |
| 80x.indel | NeuSomatic | 0.865846 | 0.708816 | 0.779501 |
| 80x.indel | VarScan | 0.756308 | 0.102085 | 0.179889 |
| 200x.indel | Strelka2 | 0.913846 | 0.757267 | 0.828221 |
| 200x.indel | Mutect2 | 0.889846 | 0.522399 | 0.658320 |
| 200x.indel | VarDict | 0.960000 | 0.049378 | 0.093925 |
| 200x.indel | RabbitVar | 0.881846 | 0.884022 | 0.882933 |
| 200x.indel | NeuSomatic | 0.908923 | 0.776143 | 0.837302 |
| 200x.indel | VarScan | 0.782769 | 0.074595 | 0.136210 |
| 300x.indel | Strelka2 | 0.913231 | 0.757916 | 0.828356 |
| 300x.indel | Mutect2 | 0.868923 | 0.513455 | 0.645486 |
| 300x.indel | VarDict | 0.960000 | 0.042953 | 0.082226 |
| 300x.indel | RabbitVar | 0.886154 | 0.878585 | 0.882353 |
| 300x.indel | NeuSomatic | 0.909538 | 0.786170 | 0.843367 |
| 300x.indel | VarScan | 0.788308 | 0.082693 | 0.149685 |

Table T7: Runtime(m) of different callers at different depths and the speedup of RabbitVar for the callers

| Depth | RabbitVar | Strelka2 | speedup | Mutect2 | speedup | VarDict | speedup | NeuSomatic | speedup | VarScan | speedup |
|---|---|---|---|---|---|---|---|---|---|---|---|
| High confidence regions | | | | | | | | | | | |
| 10x | 6 | 6 | 1.00 | 34 | 5.67 | 31 | 5.17 | 802 | 133.67 | 49 | 8.17 |
| 30x | 10 | 9 | 0.90 | 37 | 3.70 | 44 | 4.40 | 465 | 46.50 | 115 | 11.50 |
| 50x | 14 | 13 | 0.93 | 49 | 3.50 | 58 | 4.14 | 313 | 22.36 | 179 | 12.79 |
| 80x | 19 | 20 | 1.05 | 75 | 3.95 | 79 | 4.16 | 352 | 18.53 | 316 | 16.63 |
| 200x | 41 | 51 | 1.24 | 229 | 5.59 | 176 | 4.29 | 595 | 14.51 | 1147 | 27.98 |
| 300x | 59 | 76 | 1.29 | 403 | 6.83 | 264 | 4.47 | 768 | 13.02 | 4753 | 80.56 |
| All callable regions | | | | | | | | | | | |
| 10x | 10 | 11 | 1.10 | 45 | 4.50 | 52 | 5.20 | 996 | 99.60 | 59 | 5.90 |
| 30x | 13 | 52 | 4.00 | 57 | 4.38 | 158 | 12.15 | 580 | 44.62 | 142 | 10.92 |
| 50x | 16 | 81 | 5.06 | 76 | 4.75 | 259 | 16.19 | 391 | 24.44 | 222 | 13.88 |
| 80x | 21 | 111 | 5.29 | 110 | 5.24 | 362 | 17.24 | 422 | 20.10 | 352 | 16.76 |
| 200x | 43 | 155 | 3.60 | 306 | 7.12 | 781 | 18.16 | 661 | 15.37 | 1510 | 35.12 |
| 300x | 61 | 248 | 4.07 | 512 | 8.39 | 1051 | 17.23 | 934 | 15.31 | 5942 | 97.41 |

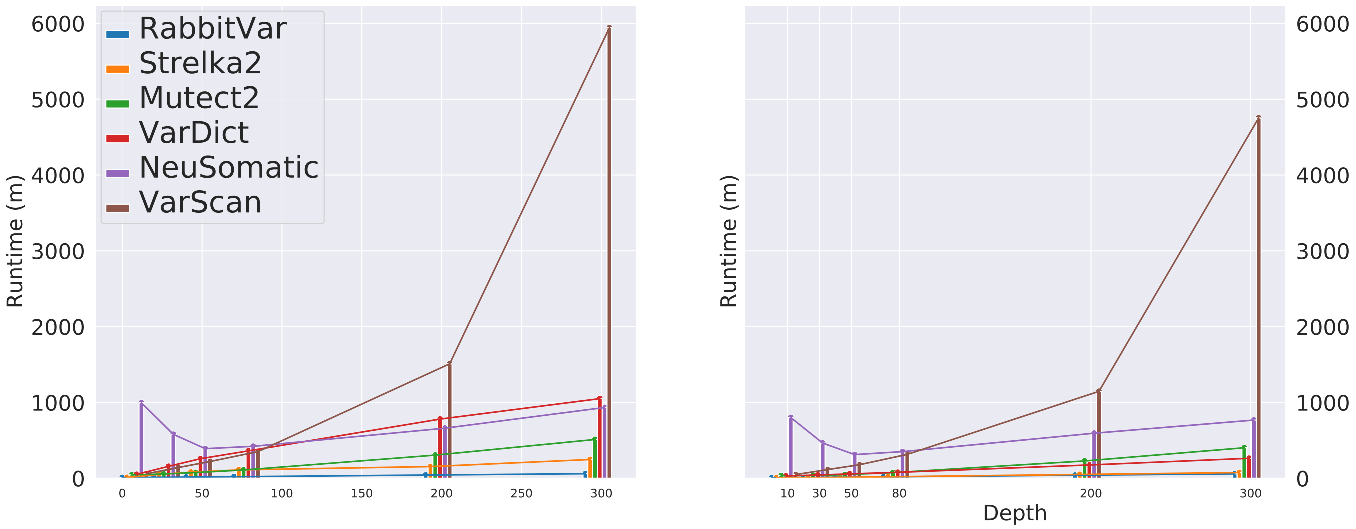


Figure S4: Runtime of different depth.

#### 1.4 Thread scalability of RabbitVar

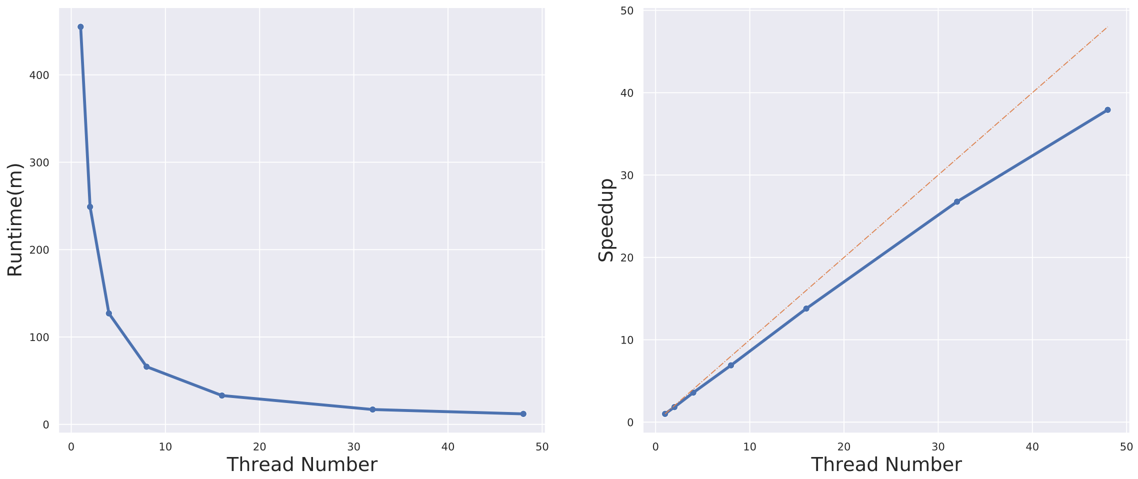


Figure S5: Runtime and thread scalability of RabbitVar on different thread number

#### 1.5 Sensitivity of different variant allele frequency on different callers

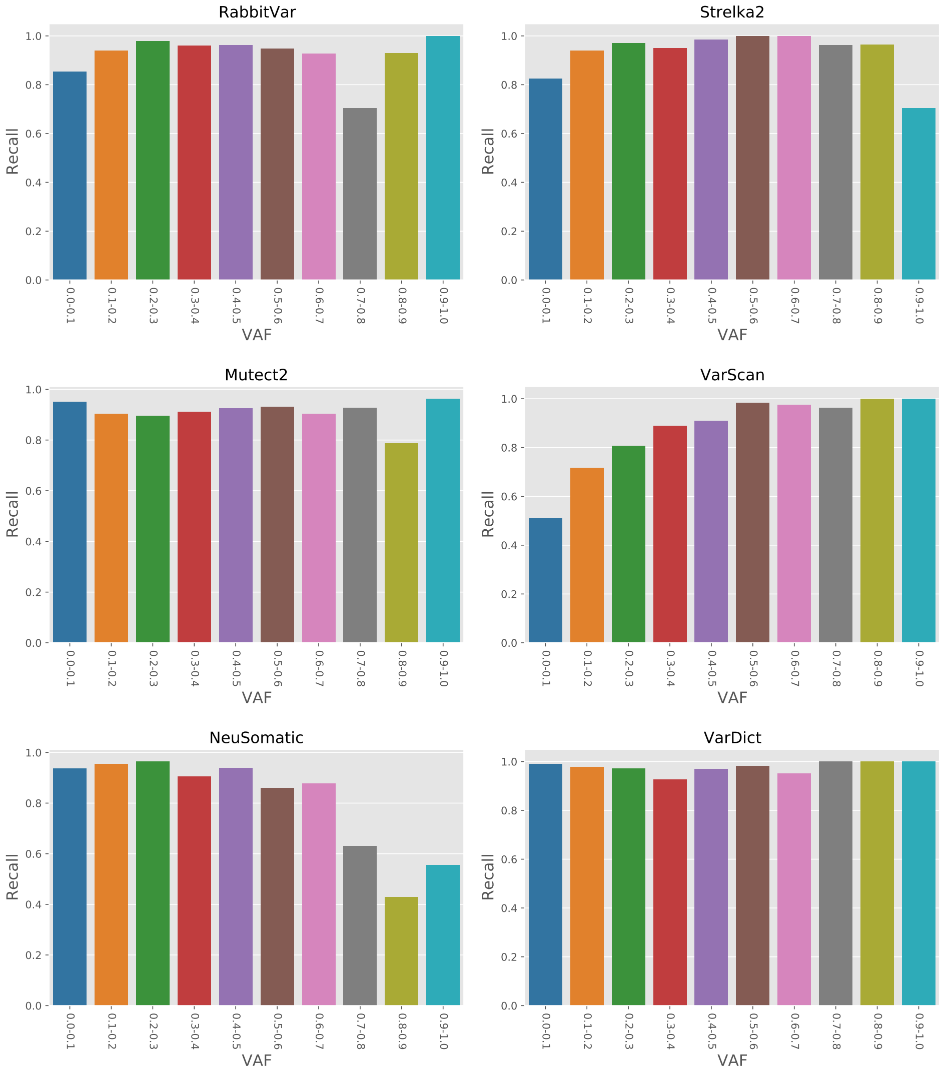


Figure S6: The indel recall evaluation on different VAF. We use the 'TVAF' field of the high-confidence truth set provided by the SEQC-II project. It shows that VarDict has the highest recall value among all callers, but the precision is very low. The experiment data is WGS_NV_1 data.

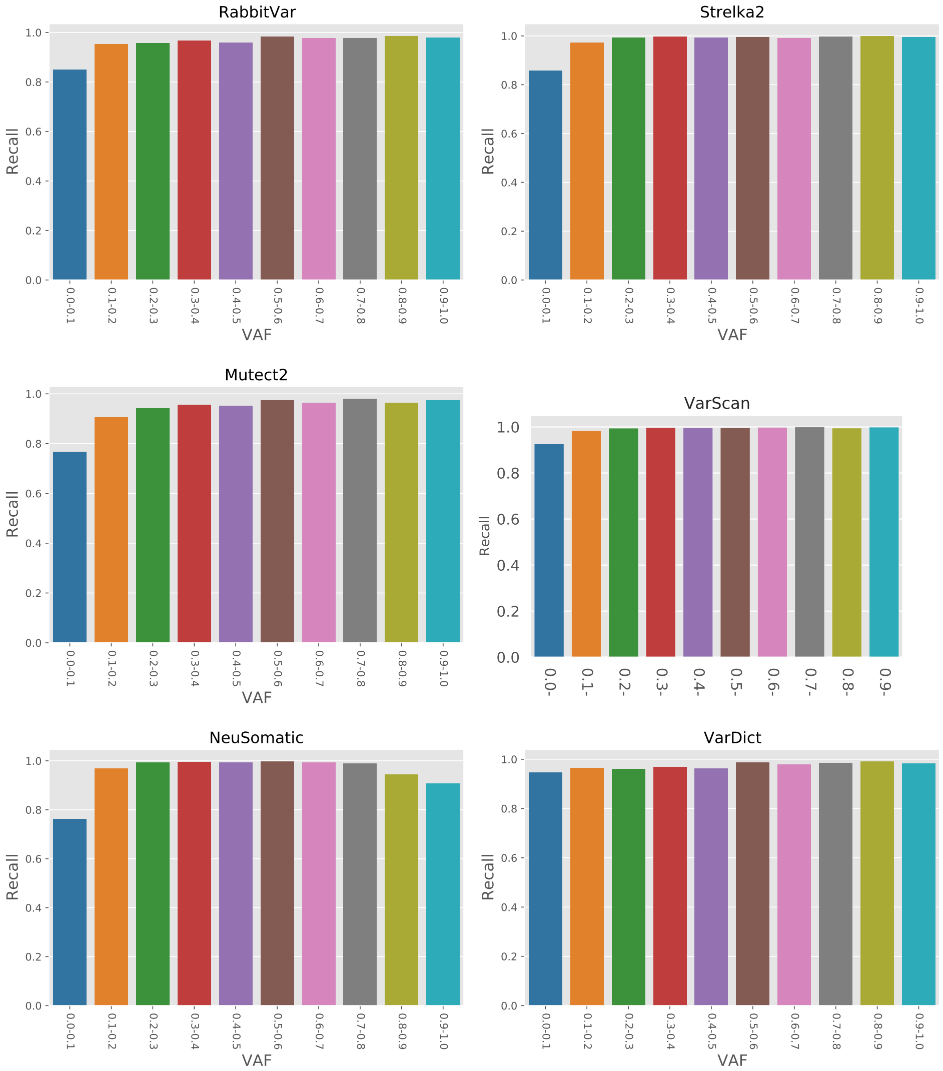


Figure S7: The SNV recall evaluation on different VAF. We use the 'TVAF' field of the high-confidence truth set provided by the SEQC-II project. It shows that VarDict has the highest recall value among all callers, but the precision is very low. The experiment data is WGS_NV_1 data.

##### 1.6 NA24631_PACA

Table T8: SNV variant calling result evaluation

| Caller | Recall | Precision | F1-score |
|---|---|---|---|
| RabbitVar | 0.8340 | 0.9780 | 0.9003 |
| Strelka2 | 0.8999 | 0.9214 | 0.9105 |
| Mutect2 | 0.8809 | 0.9286 | 0.9041 |
| NeuSomatic | 0.8355 | 0.9903 | 0.9063 |
| VarDict | 0.9429 | 0.0750 | 0.1390 |
| VarScan | 0.9443 | 0.0024 | 0.0049 |

Table T9: indel variant calling result evaluation

| Caller | Recall | Precision | F1-score |
|---|---|---|---|
| RabbitVar | 0.7609 | 0.9652 | 0.8510 |
| Strelka2 | 0.8468 | 0.9572 | 0.8986 |
| Mutect2 | 0.7630 | 0.8510 | 0.8046 |
| NeuSomatic | 0.6399 | 0.9788 | 0.7738 |
| VarDict | 0.9576 | 0.0441 | 0.0843 |
| VarScan | 0.9285 | 0.0237 | 0.0462 |

#### 2 Allel Frequency Distribution

#### 3 Feature Description

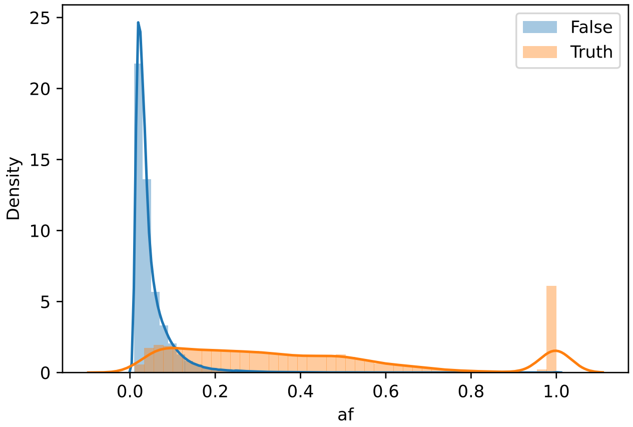


(a) SNV distribution

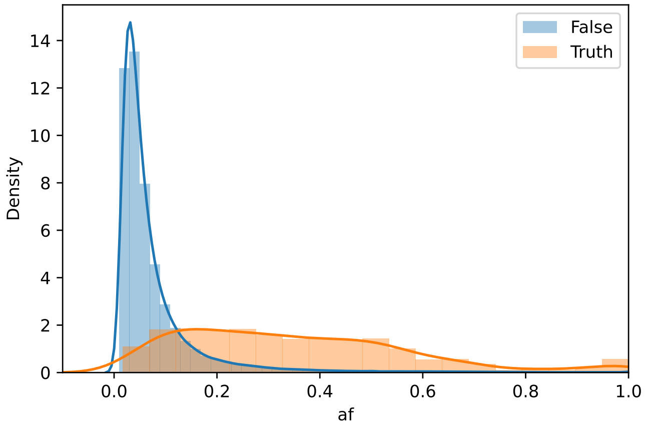


(b) INDEL distribution

Figure S8: Distribution of truth variant and false positive variants. The shown dataset is the training dataset created based on FD2 using BamSurgeon. We found other training and testing data have a similar distribution.

Table T10: Description of selected features

| Feature | Description |
|---|---|
| totalPosCov | total coverage |
| posCoverage | variant coverage |
| refFwCov | forward coverage of reference |
| refRvCov | reverse coverage of reference |
| varsFwCount | forward coverage of variant |
| varsRvCount | reverse coverage of variant |
| frequency | Allele frequency |
| meanPosition | mean position in read |
| pstd | flag for read position standard deviation (1 if the variant is covered by at least 2 read segments with different positions, otherwise 0). |
| pvalue | the pvalue from Fisher Test |
| meanQuality | mean base quality |
| ratio | odd ratio from Fisher Test |
| mapq | mean mapping quality |
| qratio | ratio of high-quality reads to low-quality reads |
| higreq | variant frequency for high-quality reads |
| shift3 | number of bases to be shifted to 3 prime for deletions due to alternative alignment |
| msi | microsatellite > 1 indicates MSI |
| msint | microsatellite unit length in bp |
| nm | average number of mismatches for reads containing the variant |
| hicnt | number of high-quality reads with the variant |
| hicov | position coverage by high quality reads |
| TumorNormalOddRatio | the logarithm of the ratio of frequencies of variants in tumor and normal.TumorNormalOddratio $= log(\frac{F_{tumor}}{F_{normal}})$ |

### 4 Scripts

#### 4.1 Runing RabbitVar

Users can run RabbitVar with two separate steps (candidate-finding and filtering):

```
time /PATH_TO_RABBITVAR/bin/RabbitVar \
  -i "/PATH_TO_BED/genome.bed" \
  -G "/PATH_TO_HG38/hg38.fa" \
  -N "${TUMOR_SAMPLE_NAME}|${NORMAL_SAMPLE_NAME}" \
  -b "${TUMOR_SAM}|${NORMAL_SAM}" \
  -c 1 -S 2 -E 3 -g 4 \
  -f 0.01 \
  --auto_resize --fisher --th ‘nproc‘ \
  --out ${TUMOR_SAMPLE_NAME}.txt
```

Then users could filter the candidate variants with:

```
python PATH_TO_RABBITVAR/XGBoost/call_xgboost.py \
  --in_file  ${TUMOR_SAMPLE_NAME}.txt \
  --model ${MODEL} \
  --scale "0.2:0.9:0.2"  \
  --var_type ${VTYPE} \
  --out_file ./${OUT}
```

Or just use the python wrapper:

```
PATH_TO_RABBITVAR/run_rabbitvar.py \
  -i "/PATH_TO_BED/genome.bed" \
  -G "/PATH_TO_HG38/hg38.fa" \
  -N "${TUMOR_SAMPLE_NAME}|${NORMAL_SAMPLE_NAME}" \
  -b "${TUMOR_SAM}|${NORMAL_SAM}" \
  -c 1 -S 2 -E 3 -g 4 \
  -f 0.01 --auto_resize --fisher --th ‘nproc‘ \
 --indelmod ./XGBoost/models/INDEL_model.pkl \
 --snvmod ./XGBoost/models/SNV_model.pkl \
 --vcf variants.vcf
```

RabbitVar support both two-segmented scale filtering strategy, for example, a parameter with *"–scale 0.2:0.9:0.2"* means when VAF $\leq$ 0.2, the predict probability (*proba*) $\leq$ 0.9 will be filtered out, and when VAF > 0.2 the *proba* $\leq$ 0.2 will be filtered out. Or a uniform scale filtering strategy, for example, *"–scale 0.5"* mean for all VAF, the variant will be filtered out when *proba* $\leq$ 0.5. The default value is "0.2:0.9:0.2".

#### 4.2 Som.py script

```
export PYTHONPATH="${BCBIO_PYTHON2.7_PATH}/site-
    packages":${PYTHONPATH}
  python ${BCBIO_PATH}/tools/bin/som.py \
  ${TRUTH} \
  ${VCF} \
  -r ${BCBIO_PATH}/genomes/Hsapiens/hg38/seq/hg38.fa \
  -R High-Confidence_Regions_v1.2.bed \
  -o ./${OUT}_${VTYPE}_sompy \
```

###### 4.2.1 Running Strelka2, Mutect2, VarDict, VarScan with bcbio-nextgen

We used the following configuration (.yaml) file:

```
details:
- algorithm:
    recalibrate: false
    mark_duplicates: false
    min_allele_fraction: 1
    variantcaller:
      somatic: [strelka2, mutect2, vardict, varscan]
  analysis: variant2
  description: TUMOR_SAMPLE
  files: TUMOR_FILE
  genome_build: hg38
  metadata:
    batch: HCC1395
    phenotype: tumor
- algorithm:
    recalibrate: false
    mark_duplicates: false
    min_allele_fraction: 1
    variantcaller:
      somatic: [strelka2, mutect2, vardict, varscan]
  analysis: variant2
  description: NORMAL_SAMPLE
  files: NORMAL_FILE
  genome_build: hg38
  metadata:
    batch: HCC1395
    phenotype: normal
fc_date: 'DATE'
fc_name: HCC1395_OUT_NAME
upload:
  dir: ./HCC1395_OUT_NAME
```

And then we run bcbio-nextgen with the following command:

```
bcbio_nextgen.py ${CONF_FILE}.yaml -n $(nproc)
```

###### 4.2.2 Running NeuSomatic

```
#first process
docker run -v ${DATA_DIR}:/mnt -v ${OUTPUT}:/output
   -v ${REF_DIR}:/ref -u $UID --memory 200G
   msahraeian/neusomatic /bin/bash -c \
"python /opt/neusomatic/neusomatic/python/preprocess
   .py \
   --mode call \
   --reference /ref/GRCh38.d1.vd1.fa \
   --region_bed /mnt/High-Confidence_Regions_v1.2.
      bed \
   --tumor_bam ${TUMOR_SAM} \
   --normal_bam ${NORMAL_SAM} \
   --work /output/work_call \
   --min_mapq 10 \
   --num_threads 64 \
   --scan_alignments_binary /opt/neusomatic/
      neusomatic/bin/scan_alignments"
echo "preporcessing done!"
time docker run --ipc=host -v ${DATA_DIR}:/mnt -v ${
   OUTPUT}:/output -v ${REF_DIR}:/ref -u $UID --
   memory 200G  msahraeian/neusomatic /bin/bash -c \
"CUDA_VISIBLE_DEVICES=0,1 python /opt/neusomatic/
   neusomatic/python/call.py \
   --candidates_tsv /output/work_call/dataset/*/
      candidates*.tsv \
   --reference /ref/GRCh38.d1.vd1.fa \
   --out  /output/${TUMOR_SAMPLE_NAME} \
   --checkpoint /opt/neusomatic/neusomatic/models/
      NeuSomatic_v0.1.4_standalone_SEQC-WGS-Spike.
      pth \
   --num_threads 64 \
   --batch_size 64 "
echo "porcessing done!"
time docker run -v ${DATA_DIR}:/mnt -v ${OUTPUT}:/
   output -v ${REF_DIR}:/ref -u $UID --memory 200G
   msahraeian/neusomatic /bin/bash -c \
"python /opt/neusomatic/neusomatic/python/
   postprocess.py \
   --reference /ref/GRCh38.d1.vd1.fa \
   --tumor_bam ${TUMOR_SAM} \
```

```
    --pred_vcf /output/${TUMOR_SAMPLE_NAME}/pred.vcf
       \
    --candidates_vcf /output/work_call/work_tumor/
       filtered_candidates.vcf \
    --output_vcf /output/post/NeuSomatic.vcf \
    --work /output/post "
  echo "done!"
```

Note that the docker image of NeuSomatic does not support calling with GPU. When evaluating the running time, we build NeuSomatic locally and run NeuSomatic with two NVIDIA GTX 3090 GPUs on the calling step. In other experiments, we just use the docker version of NeuSomatic with the script listed above.

###### 4.2.3 Get running time

```
works=(`ls -d ./SPP*/`)
for work in ${works[@]}
do
 LOG_BASE="./${work}/log"
 CMD_LOG_FILE="${LOG_BASE}/bcbio-nextgen-commands.log"
 LOG_FILE="${LOG_BASE}/bcbio-nextgen.log"
 VC_START=`grep -n "Timing: variant calling" ${
    LOG_FILE}`
 SK2_START=`grep -n "runWorkflow.py -m local -j 1 --
    quiet" \\
           ${CMD_LOG_FILE} | head -n 1`
 MT2_START=`grep -n "Mutect2 --annotation
    ClippingRankSumTest \\
           --annotation DepthPerSampleHC" \\
           ${CMD_LOG_FILE} | head -n 1`
 VDT_START=`grep -n "vardict-java -G ${PATH-TO-
    REFERENCE}" \\
           ${CMD_LOG_FILE} | head -n 1`
 VC_END=`grep "Timing: joint squaring off" ${LOG_FILE
    }`

 t0=`echo $VC_START  | grep -oP '\[.*Z\]'`
 t1=`echo $SK2_START | grep -oP '\[.*Z\]'`
 t2=`echo $MT2_START | grep -oP '\[.*Z\]'`
 t3=`echo $VDT_START | grep -oP '\[.*Z\]'`
 te=`echo $VC_END | grep -oP '\[.*Z\]'`

 t0=${t0//[\[,\],T,Z]/ }
 t1=${t1//[\[,\],T,Z]/ }
 t2=${t2//[\[,\],T,Z]/ }
```

```
 t3=${t3//[\[,\],T,Z]/ }
 te=${te//[\[,\],T,Z]/ }

 sktime=$(((`date -d "$t2" '+%s'` - `date -d "$t0" '+%
    s'`) / 60))
 mutect=$(((`date -d "$t3" '+%s'` - `date -d "$t2" '+%
    s'`) / 60))
 vardict=$(((`date -d "$te" '+%s'` - `date -d "$t3" '
    +%s'`) / 60))
 echo "Strelka2 time: ${sktime}m, Mutect2 time: \\
      ${mutect}m, VarDict time: ${vardict}m"
done
```
